# Non-destructive label-free automated identification of bacterial colonies at the species level directly on agar media using digital holography and convolutional neural network algorithms

**DOI:** 10.1101/2025.08.14.670331

**Authors:** Prisca Perlemoine, Jordan Belissard, Bruno Burtschell, Nassim Halli, Luc Martin, Camille Brunet, Maxime Gougis, Patrick Schiavone, Yvan Caspar

**Author notes:** Correspondence: Dr Yvan CASPAR, Laboratoire de Bactériologie-Hygiène Hospitalière, Institut de Biologie et Pathologie, Centre Hospitalier Universitaire Grenoble Alpes, CS10217 38043 GRENOBLE Cedex 9, FRANCE.

## Abstract

**Objectives:** this study aimed to develop a fully automated, non-destructive and label-free identification method of bacterial colonies, directly on agar plates, using a combination of digital holography and artificial intelligence and to evaluate its performances.

**Method:** high-resolution holographic images of individual colonies on translucent brain-heart agar plates were taken every 30 minutes throughout an 18-hour incubation period (530 MPx for the full plate) using a large field 1x magnification system, a partially coherent LED light source and a high-resolution CMOS sensor. A database containing 49 490 digital holograms of individual colonies from 276 clinical strains belonging to ten of the most prevalent pathogenic bacterial species was used to train the convolutional neural network (CNN). Improvement in the accuracy of the prediction from the CNN algorithms was achieved using the information at different phylogenetic levels.

**Results:** the performance of the BAIO-DX solution was assessed on 232 strains belonging to the 10 species used to train the algorithms but also on 64 strains from 8 species not included in the training database. For the species included in the training dataset, this new method identified 86.6% of the strains at the species level with a positive-percent agreement of 96.5%. An additional 48% of the strains not identified at the species level could be identified at the genus level thanks to the phylogenetic interpretation of the results.

**Conclusions:** these first results validate this approach as a candidate to obtain a fully automated non-destructive and label-free solution for bacterial identification in clinical microbiology laboratories.

**IMPORTANCE STATEMENT:** Identification of pathogenic bacteria by culture-based methods are typically performed using MALDI-TOF mass spectrometry or biochemical systems. While automation and interpretive algorithms based on agar plate imaging and artificial intelligence (AI) has reduced manual steps, bacterial identification is still labor-intensive. Here we developed a fully automated, non-destructive and label-free identification method of bacterial colonies at the species level, directly on agar plates, using a combination of digital holography and convolutional neural network algorithms. After training the system with 276 strains belonging to ten of the most frequent pathogenic bacterial species, the BAIO-DX solution was able to identify 86.6% of new strains from these 10 species with a positive-percent agreement of 96.5%. These thorough proof of concept shows that imaging methods coupled to AI algorithms are promising to reach a fully automated identification of a significant proportion of pathogenic bacteria and has potential to enhance diagnostic workflows in clinical microbiology.

## INTRODUCTION

While clinical bacteriology has been very manual and labor intensive for decades, culture-based methods have been revolutionized by the introduction of MALDI-TOF mass spectrometry and Total Laboratory Automation (TLA) systems [1]. Through robotics and software developments, TLA systems associate inoculation/streaking unit(s), incubation system(s) and high-resolution imaging of the colonies on the agar plates. More recently, artificial intelligence (AI) and interpretive algorithm software have been introduced in the workflow, facilitating the reading of urine samples or of multidrug-resistant bacteria screenings. They also allow the sorting of negative agar plates in a few minutes and for automated colony count and colony color analysis on positive agar plates, providing the possibility to detect polymicrobial samples. It has facilitated 24/7 operations, replaced batch reading of agar plates by more continuous readings, thus reducing turnaround times and providing technologist time savings that can be redirected to other diagnosis methods [2–4].

However, these technologies are not affordable for all laboratories, especially in low- and middle-income countries. Even in high income countries, despite the development of modules that allow to automatize picking and deposition of bacterial colonies, the colony transfer on MALDI-TOF target mainly remains a manual task [2]. Moreover, reading of the digital pictures of the plates requires trained staff that may not be available 24/7, while early identification of bacteria in samples like positive blood cultures (PBC) is of crucial interest as early result increase favorable outcomes [1]. The later times have also seen a shortage of trained technologists in many public or private laboratories and the difficulty in recruiting staff for evening and night shifts [1,5]. In this context, an affordable and fully automated method able to identify bacterial species directly on agar plate without any manual intervention would be of great value.

Very high-resolution holographic image (hologram) encodes the three-dimensional object complex transmittance into a 2D image. Importantly, phase objects such as bacterial colonies can be imaged with good contrast. This study aims at presenting a thorough proof of concept of a new rapid method for automated, non-destructive and label-free identification of bacterial colony directly on agar plates, at the species level, by digital in-line holography (DIH) combined convolutional neural networks (CNN) algorithms. We hypothesized that analysis of holograms of growing bacterial colonies by CNN inference processes could allow bacterial identification at the species level. Building on previous works, we developed a technology and an automaton able to process standard 90 mm brain heart infusion (BHI) agar plates [6]. The main objective of this study was to construct a first database using the BAIO-DX prototype, to develop the AI algorithms and to assess the performance of this method using clinical strains.

## MATERIALS AND METHODS

### Clinical samples, bacterial strains and sample preparation

We used 104 monomicrobial positive blood cultures (PBC) collected prospectively between August 1^st^ 2022 to January 31^st^ 2023 and 469 strains from 19 different species isolated from various clinical samples and conserved in the collection of the bacteriology laboratory of Grenoble Alpes University Hospital. Written consent for participation was not required for this study, in accordance with national legislation and institutional requirements. As part of standard of care (SoC) diagnosis procedure of bacteremia an aliquot of the PBC bottles (BD BACTEC™ aerobic/F, Becton Dickinson, Pont de Claix, France) was transferred into a dry tube (BD Vacutainer) and diluted 1/50 in sterile water tube (Saline Solution, Becton Dickinson). If bacterial identification following SoC protocol as previously described, showed a monomicrobial bacteremia, 100 µl of the bacterial suspension were plated on brain heart infusion agar plates (BHI agar, Becton Dickinson) using TLA (BD Kiestra™ Inoqula sample processor with the InocStreak S200 zigzag streaking pattern) [7]. Similarly, biobank strains were grown on BHI agar plates and incubated at 35 °C overnight. Then, colonies were suspended into saline solution at a turbidity of 0.5 McFarland and the suspension was diluted 1/50 in saline solution tube before plating using the same TLA process. After inoculation, BHI plates were immediately incubated in the BAIO-DX prototype (Figure 1).

**Figure 1:**
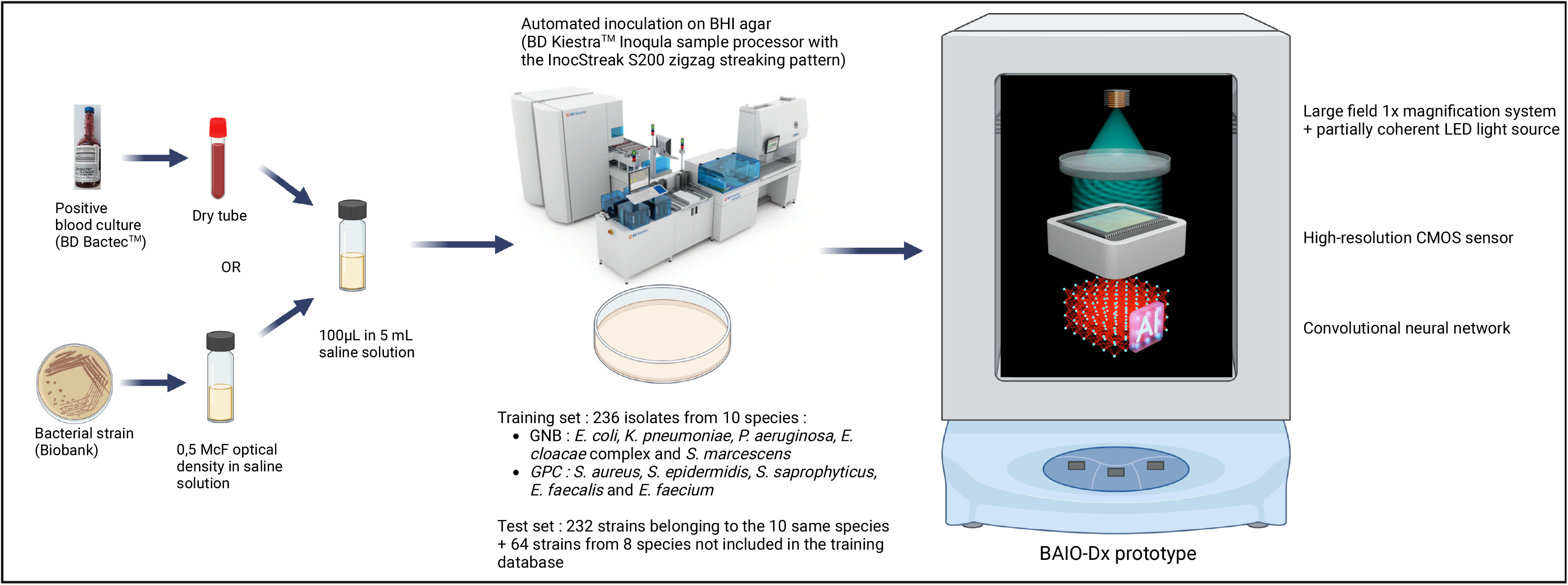
Schematic overview of the BAIO-DX digital inline holography system and study workflow. The diagram illustrates the experimental and analytical pipeline used for non-destructive, label-free bacterial identification directly on BHI agar plates. Clinical strains or monomicrobial positive blood cultures were plated and incubated within the BAIO-DX prototype, which integrates within an incubator a robotic arm and a digital in-line holography imaging system. At 30-minute intervals over an 18-hour period, high-resolution holographic images of growing colonies were acquired using a large field 1× magnification setup with a partially coherent LED light source and a high-resolution CMOS sensor. Colonies were detected and segmented, and time-lapse image stacks were processed by convolutional neural network algorithms trained on a large database of annotated holograms to predict bacterial species.

### Digital holographic imaging (DHI)

DHI was conducted using the BAIO-DX system (Figure 1). A very high-resolution (9568 × 6380 pixels) holographic image was obtained using a large field 1x magnification system, a partially coherent LED light source (ILH-ON01-DEBL-SC211-WIR200, Intelligent LED Solutions) and a high resolution monochromatic complementary metal–oxide–semiconductor (CMOS) sensor (IMX 455 from Sony; 3.76 µm pitch). The 90 mm circular agar plate was scanned over the 24×36 mm sensor and the entire plate was pictured every 30 minutes throughout an 18-hour incubation period (twelve images per time point to cover the whole plate). We adapted the workflow presented by Wang et al. with modifications allowing better automation for a higher throughput [6]. The BAIO-DX prototype has been designed to fit into a 115l Binder incubator providing enough space for the DHI system and an 18-shelf storage unit within the incubator. At every time point, a robotic arm takes each dish from its shelf, removes the cover and places the dish on the CMOS sensor for imaging and automated analysis of the agar plates.

### Data processing and AI framework for bacteria identification

The 12 holograms obtained at each time point were stitched together in order to get a final 530 MPx image of the full plate. The resulting images were aligned over time and colonies were detected using classical segmentation algorithms. For each dish, we aimed at selecting up to 200 isolated colonies. For each selected colonies, a 36-frame 125×125 pixel image stack centered on the colonies was created and used as input for the algorithms (Figure 2). Then, the images were analyzed using a CNN architecture based on a series of 3-dimensional convolutional layers combined with Spatially-Adapted Squeeze-Excitation (SASE) blocks [8–10]. We used a training set of 276 plates from 10 species. The training process was repeated 20 times to obtain 20 models. Each model was trained using a random split of 70% for training and 30% for validation.

**Figure 2:**
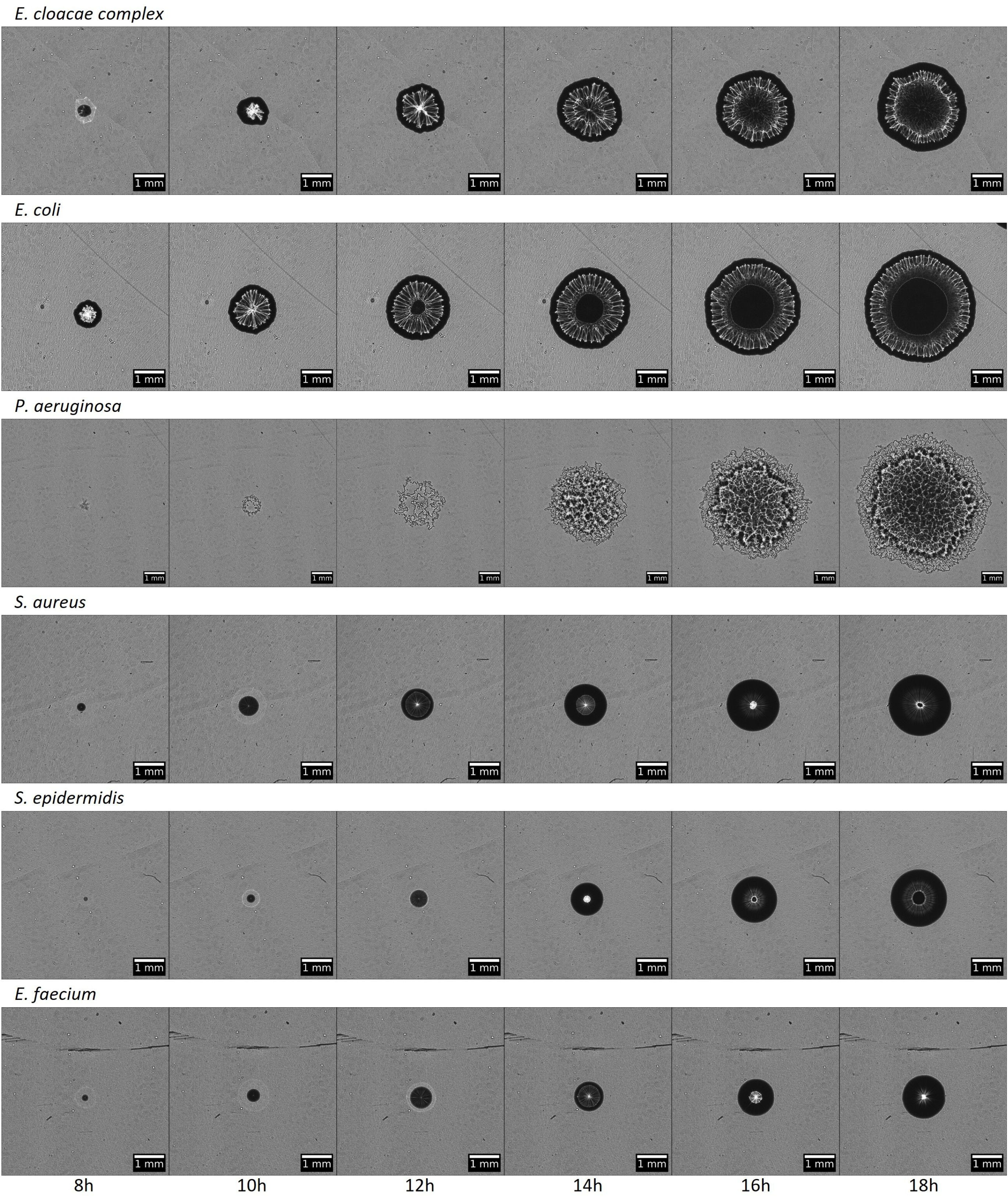
Time lapse digital holographic images of a single bacterial colony every two hours between 8h and 18h for 6 different bacterial species. Representative digital holographic images of single colonies from six different bacterial species, acquired every two hours from 8 to 18 hours of incubation on BHI agar plates. The high-resolution images capture the morphological and optical evolution of the colonies over time, highlighting species-specific growth dynamics and structural features. These time-lapse image sequences served as the input data for convolutional neural network-based species identification.

### Performance assessment of BAIO-DX system

We assessed the performance of the method to identify bacteria using another collection of 296 clinical strains and PBC samples. For each individual colony to test, a neural network inference process using all 20 models was queried and provided the species predicted with the highest score. If the confidence score was below 0.8, the identification result at the species level was rejected. Data were also classified according to a phylogenetic tree. The CNN models provided identification scores for each taxonomic level (species, genus, family, order, class, phylum, or domain). The identification prediction was performed by browsing the tree from the root, following the nodes with the highest scores until the score falls below the chosen threshold. The last node on the path with a confidence score above the chosen threshold was the result provided to the user. The level-specific thresholds were fitted in order to achieve an accuracy ≥97%. See supplemental material for additional details.

### Statistics

Identification results obtained with the BAIO-DX system were compared to SoC identification results. We calculated positive percent agreement (PPA) of the innovative method as follows: PPA (%) = 100 x number of strains matching the identification obtained by SoC method/total number of strains tested. Performance of the method is also presented as a confusion matrix of the identification results.

## RESULTS

### Non-destructive label-free automated bacterial identification at the species level

A set of 276 bacterial isolates or monomicrobial positive blood culture samples from 10 bacterial species grown on translucent BHI agar plates was used to train the BAIO-DX system (Table 1 and Figure 1). Digital imaging of the plates every 30 minutes throughout an 18-hour incubation period provided high resolution time-lapse digital holographic images of the growing bacterial colonies (Figure 2). In average, 179 +/- 42 individual colonies were analyzed per plate for a total of 49 490 individual colonies analyzed during the training process. Performance of the BAIO-DX solution was then assessed on 232 different isolates belonging to the 10 same species (Table 1). Overall, the method correctly identified the species for 201/232 (86.6%) of the strains, with a PPA of 96,5% while 31/232 (13,4%) had a score below the confidence threshold at the species level, leading to the rejection of the predicted identification. The confusion matrix of the bacterial identifications that passed the confidence threshold is presented in Figure 3. It evaluates both the accuracy of the identification results and the confusion between the predicted and the true species. The use of a phylogenetic approach and confidence thresholds at each taxonomic level (Fig S1-S3) allowed to significantly reduce misidentifications from 9,9% to 3,5% (data before rejection are presented in Figure S1), yet decreasing the percentage of strains with a valid identification at the species level.

**Table 1:**
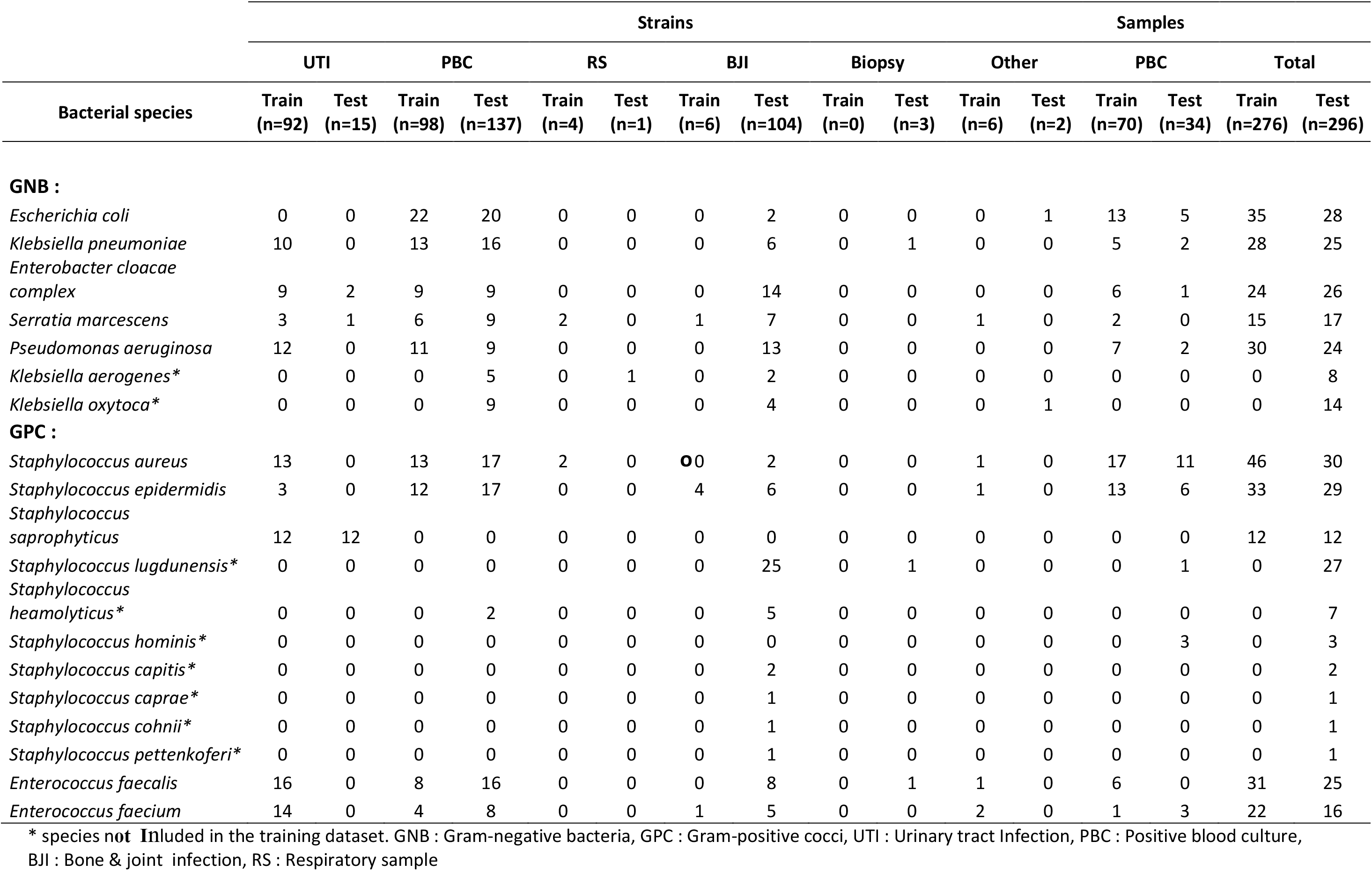
Origin and distributions of the clinical strains and clinical samples in the train and test datasets.

**Figure 3:**
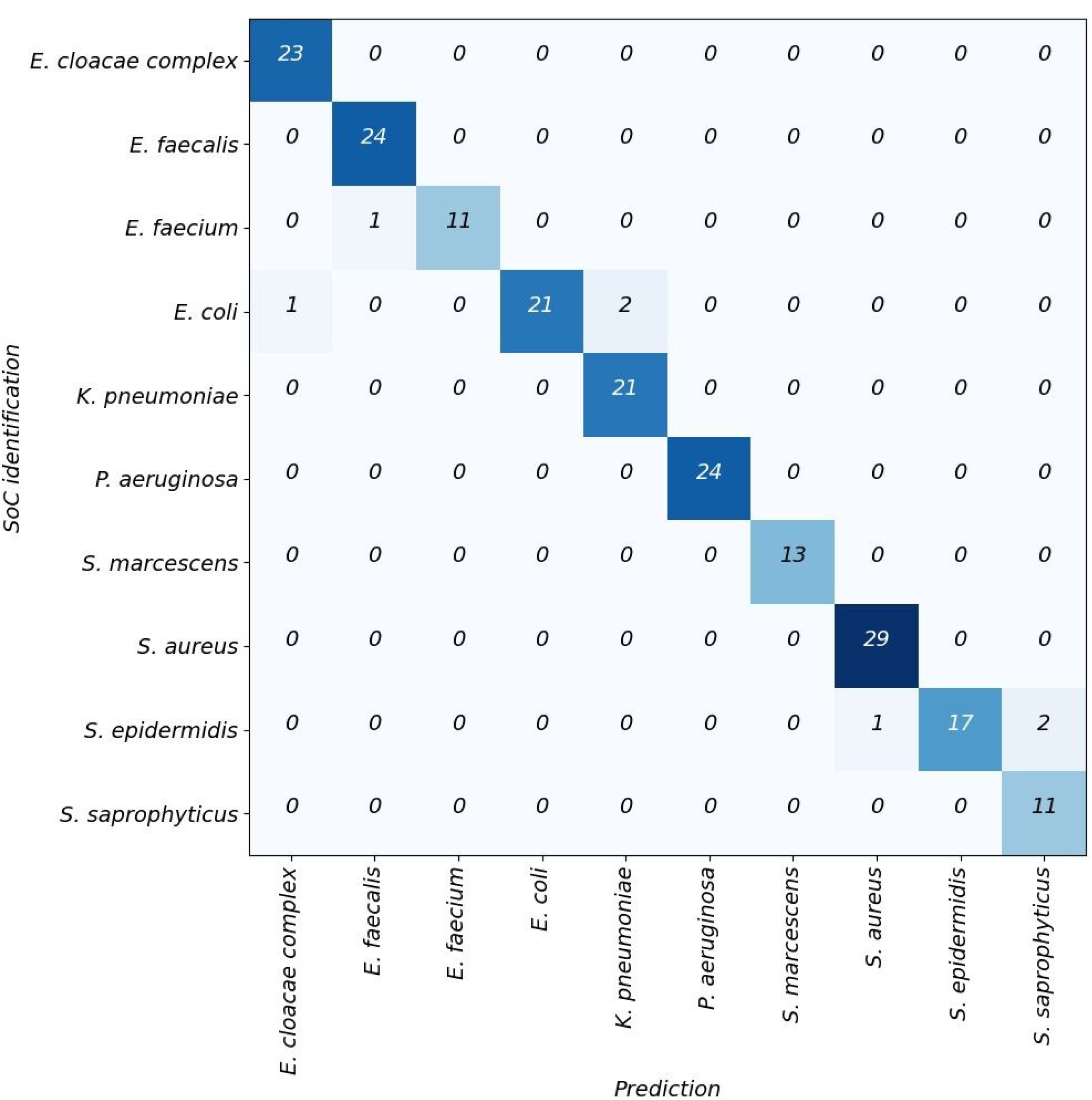
Confusion matrix of bacterial species identification using the BAIO-DX system and the phylogenetic thresholds. Confusion matrix comparing species-level predictions made by the BAIO-DX system to the reference identifications obtained by MALDI-TOF mass spectrometry for the 201 clinical isolates tested belonging to the species included in the training dataset and passing the species-level confidence threshold. Correct predictions appear along the diagonal, while off-diagonal values indicate misidentifications. The matrix illustrates the overall accuracy and the distribution of classification errors, demonstrating high concordance between the automated holography-based approach and standard-of-care (SoC) methods.

### Identification at a higher taxonomic level is possible for strains below confidence threshold at the species level

The use of a phylogenetic approach with confidence thresholds at each taxonomic levels allowed 15/31 (48%), 8/31 (26%) and 7/31 (23%) of the rejected samples at the species level to be identified at the Genus, Family or Order level respectively, with 100% PPA (Table 2).

**Table 2:**
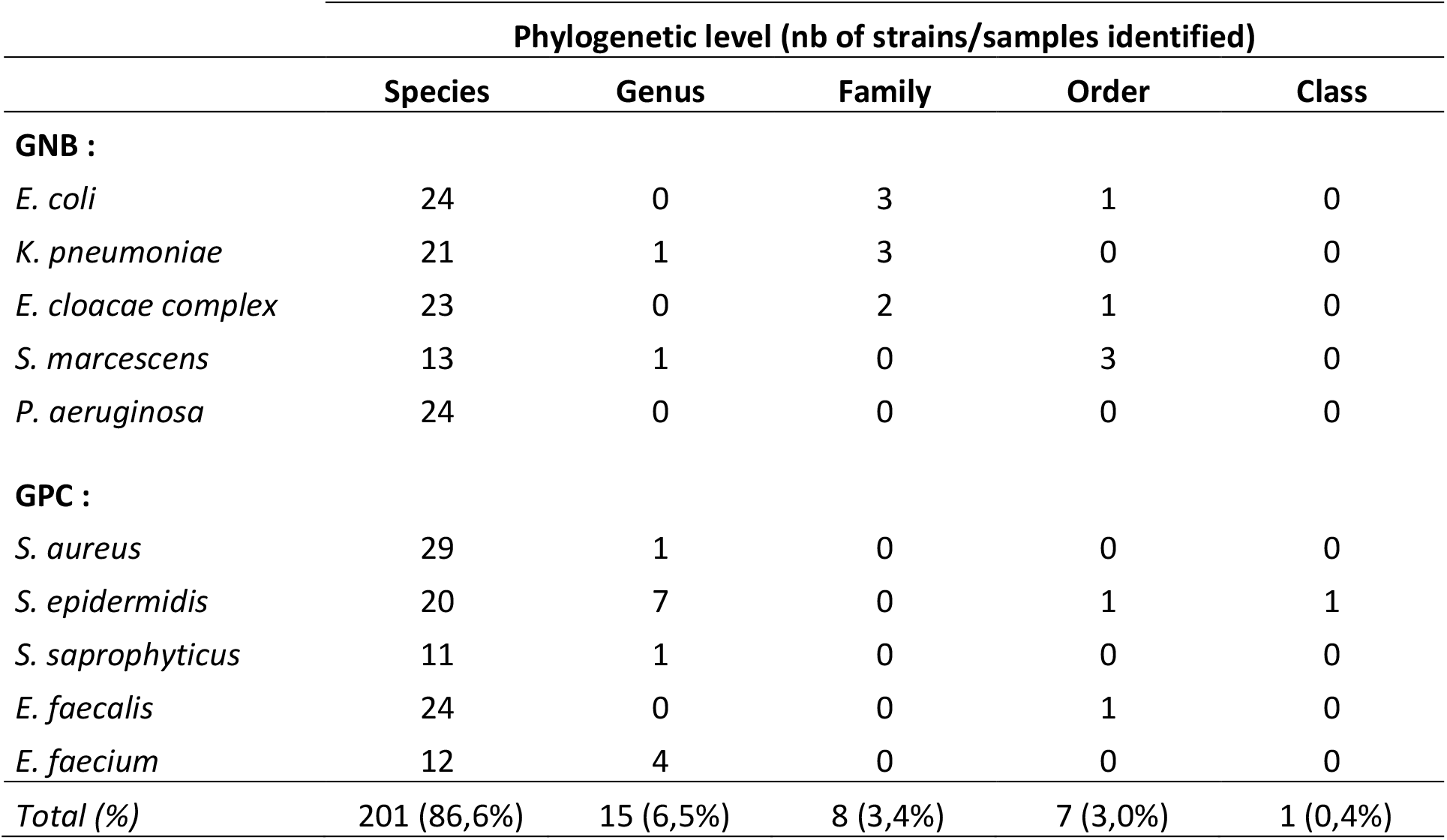
Phylogenetic level of identification provided by BAIO-DX system for the 232 isolates tested (belonging to the 10 species of the training dataset)

### Behavior of the system for species not present in the training set

Finally, we also evaluated the accuracy of this first version of the BAIO-DX solution against an “unknown” subset of strains composed of 64 isolates from 9 species not present in the training set but belonging to two bacterial genus included in the training set (42 coagulase-negative *Staphylococcus* and 22 *Klebsiella* strains) (Table1, S1 and S2). While multiclass AI systems always try to identify the closest match up among existing class, only 19/64 (29.7%) isolates were incorrectly identified at the species level thanks to the use of the phylogenetic thresholds (Table S1 and S2). Moreover 15/16 (94%), 7/7 and 15/15 strains were correctly identified at the Genus, Family or Order levels.

## DISCUSSION

In this study we elaborated and evaluated the performance of a new automated non-destructive and label-free system allowing identification of bacterial colony at the species level using digital in-line holography combined with convolutional neural network algorithms, directly on agar plates. This study shows that the first version of the BAIO-DX system, trained with a large dataset of strains from 10 species among the most commonly isolated from clinical sample (for example ≥65% of PBC isolates or ≥90% of urine samples), is able to correctly identify at the species level 86,6% of monomicrobial routine samples containing one of these 10 species with an accuracy of 96,5% [11]. Identification was below confidence threshold at the species level for 13,4% of the isolates but among those 48%, 26% and 23% could be identified at the Genus, Family or Order level respectively, with no misidentification. All strains of *S. aureus, K. pneumoniae, E. faecalis, E. cloacae complex, S. marcescens* and *S. saprophyticus* reaching the confidence threshold were correctly identified at the species level. A misidentification was observed for only 3% of the samples, for rare *E. coli, S. epidermidis* or *E. faecium* isolates. This thorough proof of concept validates that high-resolution holograms related to the morphology and the physicochemical properties of the colonies of the corresponding microorganisms (optical index, geometry, chemical bonds) have the potential to allow bacterial identification at the species level.

Such accuracy had never been reached when analyzing 10 species using this technology so far. Previous studies used either dedicated agar pads very different from agar plates used in routine microbiology or chromogenic agar media that are not developed for all species or that would lead to more costly strategies, while we selected a translucent rich BHI agar media able of culturing most pathogenic bacterial species [12,13]. Compared to previous studies, automated data acquisition of our system significantly increased the number of samples tested [6,12,14–22]. It facilitated the acquisition of a large number of samples in the database and the training set, mandatory for the CNN algorithms to provide statistically significant results. Moreover, while standard multiclass AI categorization strategies would have provided an identification at the species level for all isolates tested, the used of prediction scores and thresholds for each phylogenetic levels, improved accuracy despite providing only an identification at a higher taxonomic rank for some isolates, that may still be useful to guide antibiotic therapy or select antibiotic susceptibility testing panel for the analyzed strain.

The main limitation of the present study remains the limited number of species included in the training dataset, despite higher than in most previous studies. The technology requires translucent media. We used BHI agar plates that allow the growth of most but not all pathogenic bacterial species and evaluated only aerobic growth conditions so far. Moreover, results were interpreted at the agar plate level analyzing all isolated colonies from a monomicrobial sample. Finally, this study resulted from data from a single center with a single prototype and will require to be expanded to larger studies to validate generalizability. Our study also has several strengths. To mitigate any potential bias related to the agar plate, the data split was performed at plate level. All colonies detected within a plate included in the training set were assigned only to the training set and were not used in the test set. Using this strategy, we strived to maintain the independence and generalizability of our results by reducing the impact of dish-related factors and strains specificity which are often overlooked in similar studies [6,12,13]. Accuracy of the system may be improved (i.e. to 99%) using the same database and training set by increasing the confidence threshold but at the cost of a higher number of rejected identifications. Identification is also possible at the colony level at lower accuracy, which could allow in the future analysis of samples with complex flora. Moreover, this system requires multiple time points resulting in a very large amount of data to reach sufficient accuracy as previously observed [12,15]. However, a reduction of the acquisition frequency to 2 or 3h seems affordable with a limited impact on the identification results. Among other strengths, compared to previous reports, we also challenged our system with isolates not included in our database. As multiclass AI systems always try to identify the closest match the use of a phylogenetic approach largely reduced misidentifications.

Current bacterial identifications by MALDI-TOF MS have a rapid turnaround time, depending on the growth speed of the bacteria on agar plates but also on the availability of trained staff while this technology can deliver identification results in real time after 18h of growth without any manual intervention which may accelerate patient management. Moreover, this technology is non-destructive, label free, does not require any additional reagent and involves less expensive automation which could improve their implementation in middle or low-income countries. It seems adapted for incorporation in existing TLA systems, could help to reduce the number of different agar media inoculated for some clinical samples and may be useful to automatically choose which AST panel to perform.

In conclusion this study paves the way for a new promising non-destructive and fully automated method for bacterial identification. It still requires important development to enlarge the database and the diversity of the species in the training dataset before clinical use. Future directions may focus first on expanding the database only to species present in some specific samples like urines or on providing accurate results at an earlier time point such as 8 hours. A higher accuracy may be obtained by multimodal strategies (using Gram stain information, classical agar plate imaging from TLA systems, …) or by modifying BHI agar with selective additives that would allow to reduce the complexity of potential species growing on the media or with chromogenic compounds to add a chromogenic information to the colonies. This technology may also be useful outside clinical microbiology laboratories to identify bacteria in environmental, water or food samples or in pharmaceutical sterility controls.

## ACKNOWLEDGEMENTS

The authors would also like to thank the entire BAIO-DX team for their invaluable support and contributions to the project, as well as the staff of the Bacteriology Laboratory at Grenoble University Hospital for their assistance and expertise. Some figures were Created in BioRender. Part of the work has been presented at the French congress “Réunion interdisciplinaire de chimiothérapie anti-infectieuse”, December 18^th^ 2023.

## Contributor Roles

P.P. Y.C. M.G. and P.S. designed the study. P.P. and C.B. carried out the experiments. P.P. and Y.C. wrote the manuscript with support from M.G., J.B. and P.S.. J.B. and N.H. designed image stitching and alignment, and predictions aggregations. B.B., N.H. and L.M. designed AI framework: model architecture, training and evaluation strategy. All authors have read and agreed to the published version of the manuscript.

